# Modeling Continuous Admixture

**DOI:** 10.1101/026278

**Authors:** Ying Zhou, Hongxiang Qiu, Shuhua Xu

## Abstract

Human migration and human isolation serve as the driving forces of modern human civilization. Recent migrations of long isolated populations has resulted in genetically admixed populations. The history of population admixture is generally complex; however, understanding the admixture process is critical to both evolutionary and medical studies. Here, we utilized admixture induced linkage disequilibrium (LD) to infer occurrence of continuous admixture events, which is common for most existing admixed populations. Unlike previous studies, we expanded the typical continuous admixture model to a more general admixture scenario with isolation after a certain duration of continuous gene flow, and we demonstrated that such treatment significantly improved the accuracy of inference under complex admixture scenarios. Based on the extended models, we developed a method based on weighted LD to infer the admixture history considering continuous and complex demographic process of gene flow between populations. We evaluated the performance of the method by computer simulation and applied our method to real data analysis of a few well-known admixed populations.

## Introduction

Human migrations involve gene flow among previously isolated populations, resulting in the generation of admixed populations. In both evolutionary and medical studies of admixed populations, it is essential to understand admixture history and accurately estimate the time since population admixture because genetic architecture at both population and individual levels are determined by admixture history, especially the admixture time. However, the estimation of admixture time is largely dependent on the precision of the applied admixture models. Several methods have been developed to estimate admixture time based on the Hybrid Isolation (HI) model (Xu and Jin 2008; Price *et al.* 2009; Loh *et al.* 2013; Qin *et al.* 2015) or intermixture admixture model (IA) (Zhu *et al.* 2004), which assumes the admixed population is formed by one wave of admixture at a certain time. However, the one-wave assumption often leads to under-estimation when the progress of the true admixture cannot be well modeled by the HI model. Jin et al. have shown earlier that under the assumption of HI, the estimated time is half of the true time when the true model is a gradual admixture (GA) model (Jin *et al.* 2013).

Admixture models can be theoretically distinguished by comparing the length distribution of continuous ancestral tracts (CAT) (Gravel 2012; Jin *et al.* 2012; Ni *et al.* 2015), which refer to continuous haplotype tracts that were deviated from the same ancestral population. CAT inherently represents admixture history as it accumulates recombination events. Short CAT always indicate long admixture histories of the same admixture proportion, whereas long CAT may indicate a recent gene flow from the ancestral populations to which the CAT belong. Based on the information it provides, CAT can be used to distinguish different admixture models and estimate corresponding admixture time. However, accurately estimating the length of CAT is often very difficult.

Weighted linkage disequilibrium (LD) is an alternative tool that can be used to infer admixture (Loh *et al.* 2013; Pickrell *et al.* 2014). Previous studies have indicated that this tool is more efficient than CAT because it requires neither ancestry information inference nor haplotype phasing, which often provides false recombination information, thus decreasing the power of estimation. Weighted LD has already been used in inferring multiple-wave admixtures (Zhou *et al.* 2015). However, these methods tend to summarize the admixture into different independent waves, even if the true admixture is continuous. In our previous work (Zhou *et al.* 2015), we mathematically described weighted LD under different continuous models, allowing us to determine admixture history using these models.

In the present study, we first developed a weighted LD-based method to infer admixture with HI, GA, and continuous gene flow (CGF) models (Pfaff *et al.* 2001), (Fig 1). Both GA and CGF models assume that gene flow is a continuous process. Next, we extended the GA and CGF models to the GA-I and CGF-I models, respectively (Fig 1), which model a scenario with a continuous gene flow duration followed by a period of isolation to present. We applied our method to a number of well-known admixed population and provided information that would help better understand the admixture history of these populations.

**Fig. 1:**
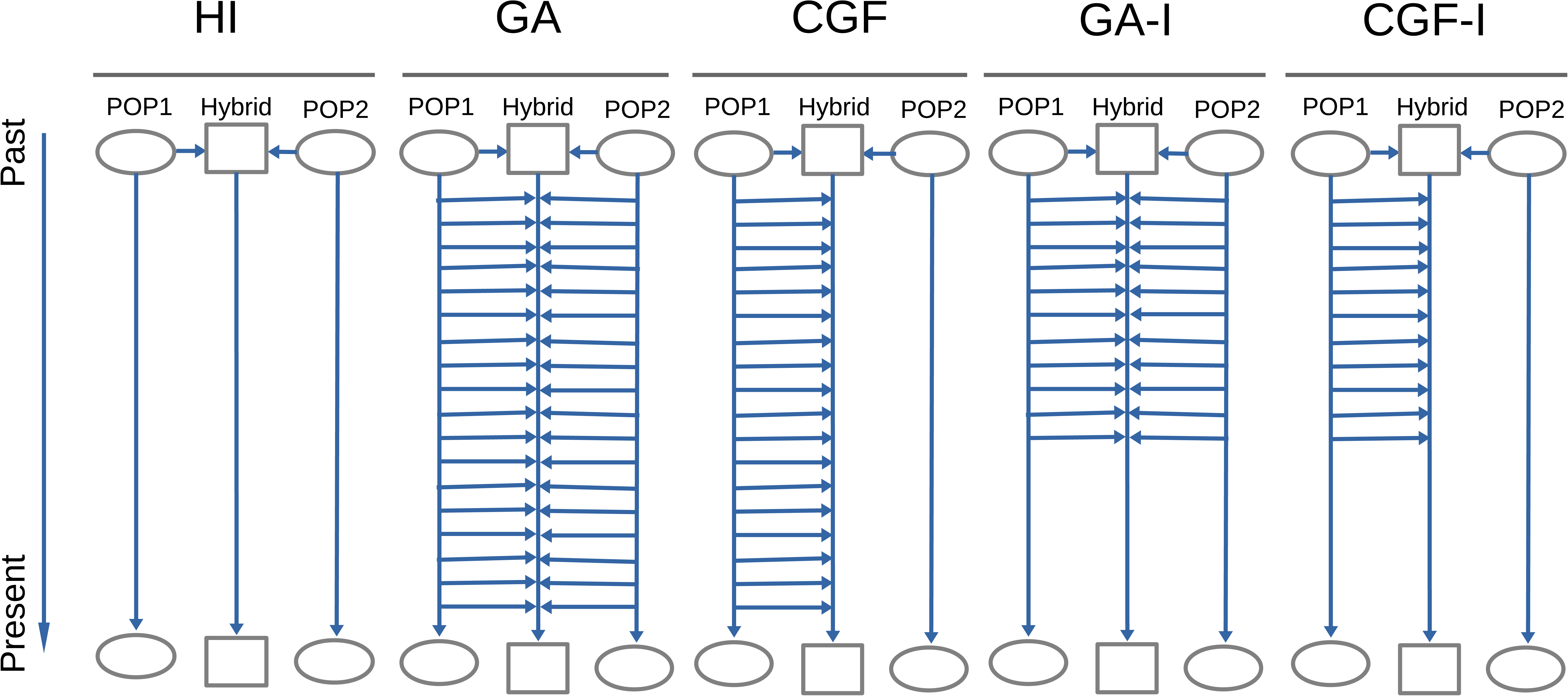
Classic admixture models (HI, GA and CGF) and the models we extended (GA-I and CGF-I). For each model, the simulated admixed population (Hybrid) is in the middle of two source populations (POP1 and POP2). Each horizontal arrow represents the direction of gene flow from the source populations to the admixed population. Once the genetic components flow into the admixed population, the admixed population randomly hybridizes with other existing components. The existence of horizontal arrows indicate gene flow from the corresponding source population.

## Materials and Methods

### Data Sets

Data for simulation and empirical analysis were obtained from two public resources: Human Genome Diversity Panel (HGDP) (Li *et al*. 2008), the International HapMap Project phase III (Frazer *et al*. 2007) and the 1000 Genomes Project (1KG) (The 1000 Genomes Project Consortium 2012). Source populations for simulations are the haplotypes from 113 Utah residents with Northern and Western European ancestries from the CEPH collection (CEU) and the 113 Africans from Yoruba (YRI).

### Inferring Admixture Histories by using the HI,GA,and CGF Models

The expectation weighted LD under two-way admixture model has been described earlier (Zhou *et al*. 2015). Following the previous notation, the expectation between two sites separated by a distance *d* (in moragan) is as follows :

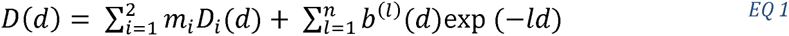

Where *b*^(*l*)^(*d*) = *c*^(*l*)^*E*((*δ*_12_(*x*)*δ*_12_(*y*))^2^ ‖*x* − *y* | = *d*); *D*(*d*) and *D*_*i*_(*d*) are the expected weighted LD of the admixed population and the source population *i*; respectively; *m*_*i*_ is the admixture proportion from the source population *i*; and *δ*_12_(*x*) is the allele frequency difference between populations 1 and 2 at site x. To eliminate the confounding effect due to background LD from the source populations, we used the quantity, *Z*(*d*), defined as the follows, to represent the admixture induced LD (ALD).

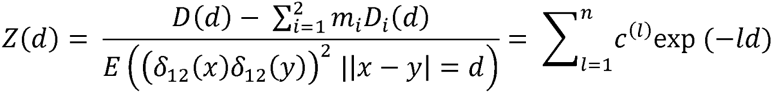

We present it in a more compact form using the inner product of two vectors as follows:

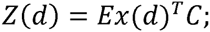

where

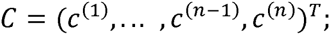

and

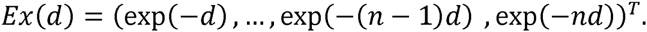

Therefore, for different admixture models where admixture began *n* generations ago, *Z*(*d*) varies in terms of the vector of coefficients of exponential functions:

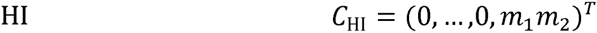

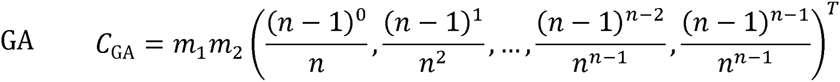

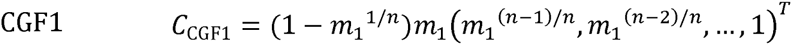

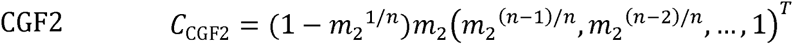

where *C*_model_ has length *n* using the HI, GA, CGF1, or CGF2 model; and *n* represents when the admixture occurred (HI) or began (GA and CGF) in terms of generations. For different models, the coefficient vectors have different patterns (Fig 2), which can be used to infer the best-fit model for a certain admixed population.

**Fig. 2:**
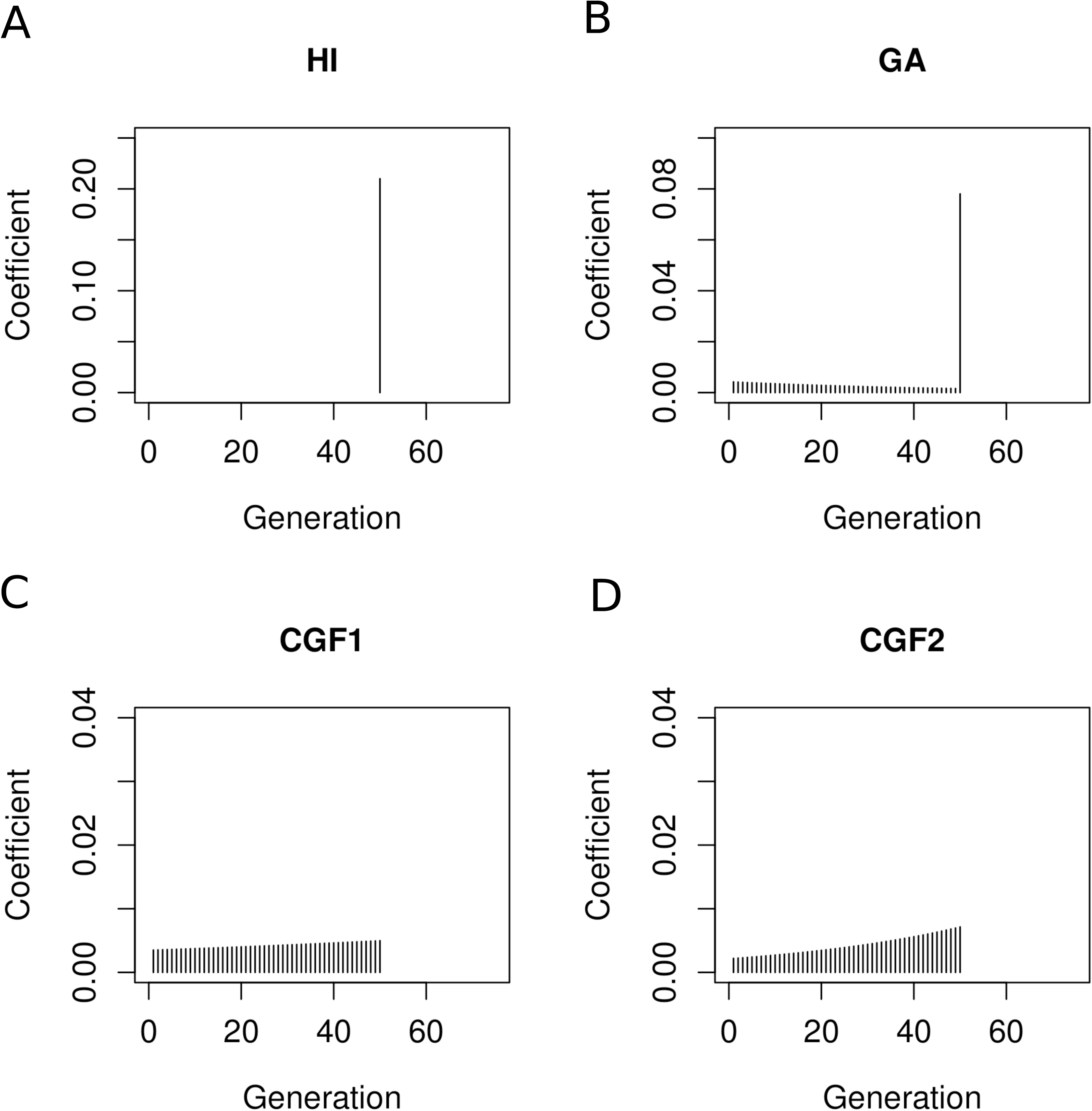
Coefficient vector of exponential functions for each model. For each admixture model, the starting time of the population admixture is 50 generations ago.

In the CGF model, CGF1 represents the admixture where source population 1 is the recipient of the gene flow from population 2, whereas CGF2 indicates source population 2 as gene flow recipient from population 1. Inference of the admixture time under different models can be regarded as minimizing the objective function as follows:

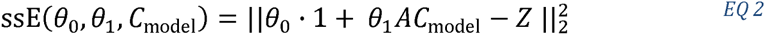

The optimization problem is therefore expressed as follows:

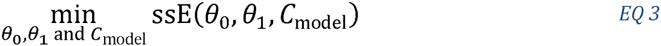

where Z= (*Z*(*d*_1_),*Z*(*d*_2_),…,*Z*(*d*_*I*_))^*T*^ is the observed ALD calculated from the single nucleotide polymorphism (SNP) data of both the parental populations and the admixed population; *θ*_0_ is a real number used to correct the population substructure; *θ*_1_ is a scalar that improves estimation robustness; **1** ∊ *R^I^* is a vector with each entry being 1; *A* is an *I* × *J* matrix with the *i*th row vector defined as *Ex*(*d*_*i*_)^*T*^, i.e., *A* = *Ex*(*d*_*1*_), *Ex*(*d*_*2*_, …, *Ex*(*d*_*I*_)^*T*^).

Next, we tried to estimate the parameters *θ*_0_,*θ*_1_, and *C*_model_, where *C*_model_ has the information of the admixture model and the related admixture time *n* (in generations). In our analysis, the value of *n* is assumed to be a positive integer; therefore, our method is to go through all possible *n* values (with a reasonable upper limit) to estimate *n* with the minimum value of the objective function. Given. *n*, we used linear regression to estimate (*θ*_0_,*θ*_1_) such that the objective function was minimized. Using this approach, the value of *n* in relation to the minimal objective function value for each model was determined, which represents the time of admixture occurrence under each model.

### Admixture Inference under HI, GA-I, and CGF-I Models

GA and CGF models assume that the admixture is strictly continuous from the beginning of admixture to present. This assumption seems too strong to be valid in empirical studies. Here, we extended the GA model and CGF model to GA-I model and CGF-I model, respectively, by considering continuous admixture followed by isolation. In this case, the admixture event lasts from *G*_start_ generations ago to. *G*_end_ generations ago. Similar to the previous case, the coefficients of exponential functions can be represented as the vector of length *G*_start_ for each model, whose first *G*_end_ – 1 entries are filled with zeros. Suppose the admixture lasted for *n* generations, then

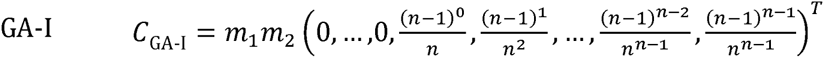

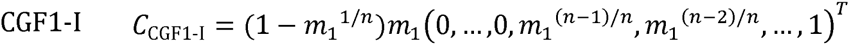

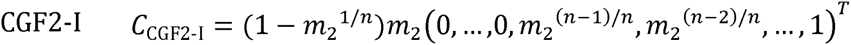

In this case, we can also try to find the parameters to minimize the objective function (EQ 2) under new models. By examining all possible pairs of (*G*_end_, *G*_start_), it is possible determine the global minimum of the objective function, although this might not be computationally efficient. Here, we used a faster algorithm (***Algorithm 1***) to determine the starting and ending time points of admixture.

Let *E* and *S* be the ending and starting time points (in generations, prior to the present) of the admixture, which we want to search for to minimize the objective function. The search starts from (*E*^0^,*S*^0^) = (1,*T_u_*), where *T_u_* is the upper bound for the beginning of the admixture event, which can be set to be a large integer to seek for a relatively ancient admixture event. In our analysis of recent admixed populations, we set *T_u_* = 500. For *k* = 1,2,…,(*E*^*k*^,*S*^*k*^) is updated from (*E*^*k*−1^,*S*^*k*−1^) by two alternative proposals. For convenience, we define

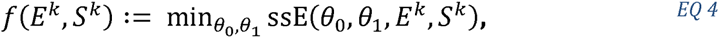

where *θ*_0_,*θ*_1_ can be determined by linear regression.

We choose the proposal that results in a smaller value for *f*. The search stops when the value of *f* with (*E*^*k*−1^,*S*^*k*−1^) is no larger than that of either proposal or *E*^*k*^ = *S*^*k*^. In this way, we can readily estimate the time interval of the admixture event (*G*_end_, *G*_start_) quickly.

#### Algorithm 1

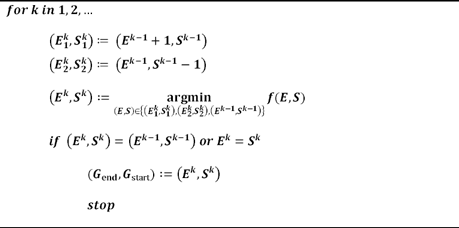

### Result evaluation

To evaluate the inference, an intuitive way is to compare empirical weighted LD with the fitted LD. Here, we use two quantities: msE and Quasi F, defined by the following:

1. Let, *e*=*θ*_0_ · 1 + *θ*_1_*AC*_model_ - *Z*. We look at 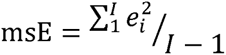 with *e*_*i*_ being the; *i*th entry of, *e*. This reflects goodness of fit and strength of background noise. A smaller msE indicates less background noise and better fit.
2. Let 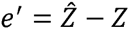, where 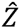 is the fitted weighted LD obtained from MALDmef, which theoretically can be regarded as the de-noised weighted LD. e′ is a vector of length *I*, with the *i*th entry denoted by 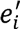. We look at the quasi-F statistic 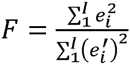. A small *F* indicates that the current fit does not significantly deviate from the previous fit.

A reliable result should have both small msE and small *F* values

### Identification of the best-fit model

For convenience of illustration, we define the **core model** as the model used to infer admixture time. When inferring admixture of a target population, HI, GA, CGF1, CGF2, GA-I, CGF1-I and CGF2-I are used as the core models for conducting inference. Because GA-I, CGF1-I and CGF2-I describe more general admixture models than GA, CGF1, and CGF2, we classified model selection into two cases: one case is to identify the best-fit model(s) among the HI, GA, CGF1, and CGF2 models, whereas the more general case is to determine the best-fit model(s) among HI, GA-I, CGF1-I and CGF2-I models. In both cases, the same strategy is adopted, which depends on the pairwise paired difference of log(msE) values associated with each core model. For an admixed population, there are 22 observed weighted LD curves (considering 22 autosomal chromosomes in an individual genome; using jackknife by leaving out one chromosome to calculate to calculated each LD curve) (Zhou *et al*. 2015). Next, we fitted the observed weighted LD curve for each core model by estimating *θ*_0_,*θ*_1_ and the time interval, which in turn allowed us to obtain the msE value associated with the optimal parameters for each weighted LD curve. Taken together, a total of 22 msE values associated with 22 LD curves were evaluated in each core model. Based on msE values calculated with different core models, the best-fit core model(s) are those with significantly small msE values. A pairwise paired t-test was conducted for the log(msE) of the four models (Table 1). When the log(msE) of HI were not significantly larger than the those of the best model, i.e., the model associated with the smallest mean of the log(msE) values, HI was selected. Otherwise the models whose log(msE) were not significantly larger than those of the best model were selected (the best model is selected as well). To control the family-wise error rate, the Holm-Bonfferoni method (Holm 1979) was used to adjust p-values, and the significance level was set to 0.05 in the present study. Here, log(msE) rather than msE were used because from our experience, log(msE) was more similar to the normal distribution.

**Table 1:**
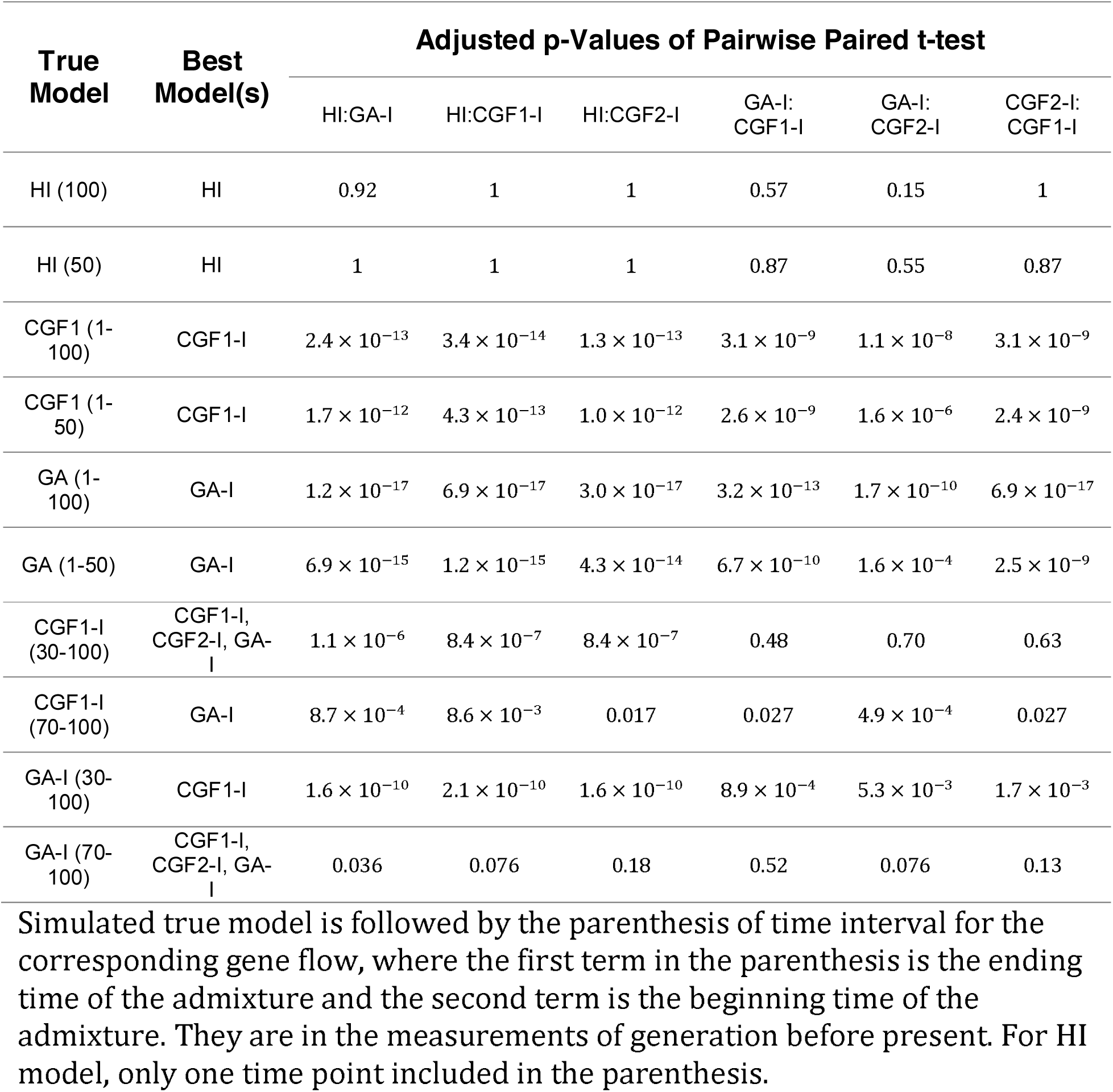
Adjusted p-values of pairwise paired t-test among core models: HI, GA-I, CGF1-I, CGF2-I.

| True Model | Best Model(s) | Adjusted p-Values of Pairwise Paired t-test |  |  |  |  |  |
| --- | --- | --- | --- | --- | --- | --- | --- |
|  |  | HI:GA-I | HI:CGF1-I | HI:CGF2-I | GA-I: CGF1-I | GA-I: CGF2-I | CGF2-I: CGF1-I |
| HI (100) | HI | 0.92 | 1 | 1 | 0.57 | 0.15 | 1 |
| HI (50) | HI | 1 | 1 | 1 | 0.87 | 0.55 | 0.87 |
| CGF1 (1-100) | CGF1-I | $2.4 \times 10^{-13}$ | $3.4 \times 10^{-14}$ | $1.3 \times 10^{-13}$ | $3.1 \times 10^{-9}$ | $1.1 \times 10^{-8}$ | $3.1 \times 10^{-9}$ |
| CGF1 (1-50) | CGF1-I | $1.7 \times 10^{-12}$ | $4.3 \times 10^{-13}$ | $1.0 \times 10^{-12}$ | $2.6 \times 10^{-9}$ | $1.6 \times 10^{-6}$ | $2.4 \times 10^{-9}$ |
| GA (1-100) | GA-I | $1.2 \times 10^{-17}$ | $6.9 \times 10^{-17}$ | $3.0 \times 10^{-17}$ | $3.2 \times 10^{-13}$ | $1.7 \times 10^{-10}$ | $6.9 \times 10^{-17}$ |
| GA (1-50) | GA-I | $6.9 \times 10^{-15}$ | $1.2 \times 10^{-15}$ | $4.3 \times 10^{-14}$ | $6.7 \times 10^{-10}$ | $1.6 \times 10^{-4}$ | $2.5 \times 10^{-9}$ |
| CGF1-I (30-100) | CGF1-I, CGF2-I, GA-I | $1.1 \times 10^{-6}$ | $8.4 \times 10^{-7}$ | $8.4 \times 10^{-7}$ | 0.48 | 0.70 | 0.63 |
| CGF1-I (70-100) | GA-I | $8.7 \times 10^{-4}$ | $8.6 \times 10^{-3}$ | 0.017 | 0.027 | $4.9 \times 10^{-4}$ | 0.027 |
| GA-I (30-100) | CGF1-I | $1.6 \times 10^{-10}$ | $2.1 \times 10^{-10}$ | $1.6 \times 10^{-10}$ | $8.9 \times 10^{-4}$ | $5.3 \times 10^{-3}$ | $1.7 \times 10^{-3}$ |
| GA-I (70-100) | CGF1-I, CGF2-I, GA-I | 0.036 | 0.076 | 0.18 | 0.52 | 0.076 | 0.13 |

Simulated true model is followed by the parenthesis of time interval for the corresponding gene flow, where the first term in the parenthesis is the ending time of the admixture and the second term is the beginning time of the admixture. They are in the measurements of generation before present. For HI model, only one time point included in the parenthesis.

## Results

### Simulation studies

Admixed populations were simulated in a forward-time way under different admixture models. For each model, simulation was performed using 10 replicates; each replicate contained 10 chromosomes with a total length of 3 Morgans. To evaluate the performance of our algorithm, we simulated admixed populations under the following conditions:

1. HI of 50 and 100 generations, designated as HI (50) and HI (100),
2. GA of 50 and 100 generations, designated as GA (1-50) and GA (1-100), respectively,
3. CGF of 50 and 100 generation, population 1 as the recipient, designated as CGF1 (1-50) and CGF1 (1-100) respectively,
4. CGF-I of a70-generation admixture followed by 30-generation isolation, and a 30-generation admixture followed by a 70-generation isolation, with population 1 as the recipient, designated as CGF1-I (30-100) and CGF1-I (70-100) respectively, and,
5. GA-I of a 70-generation admixture followed by 30-generation isolation and a 30-generation admixture followed by a70-generation isolation, designated as GA-I (30-100) and GA-I (70-100), respectively.

With simulated admixed populations, we first used the HI, GA and CGF models as core models to conduct inference (Fig S1). When the simulated model was a HI, GA, or CGF model, our method was able to accurately estimate the simulated admixture time, as well as determine the correct model, with an accuracy of 86.67%. When the simulated model was a CGF-I or GA-I model, the estimated time based on the core model HI was within the time interval of the admixture, whereas all best-fit models were HI (Table 2). This result has indicated the limitation of using the GA and CGF models in inferring admixture history.

**Table 2:** Accuracy of model detection.

| True models | Core models | Counts |  |  | Rates |  |  |
| --- | --- | --- | --- | --- | --- | --- | --- |
|  |  | Correct | Undetermined | Wrong | Correct | Undetermined | Wrong |
| HI;GA;CGF | HI;GA;CGF | 52 | 3 | 5 | 86.7% | 5.0% | 8.3% |
| GA-I; CGF-I | HI;GA;CGF | 0 | 0 | 40 | 0.0% | 0.0% | 100.0% |
| HI;GA;CGF | HI;GA-I; CGF-I | 47 | 11 | 2 | 78.3% | 18.3% | 3.4% |
| GA-I; CGF-I | HI;GA-I; CGF-I | 6 | 22 | 12 | 15.0% | 55.0% | 30.0% |

2 Here, as our method can hardly distinguish CGF1 from CGF2 model, we regard CGF1, CGF2 as the CGF model; CGF1-I and CGF2-I as the CGF-I model, which is different from GA-I and HI models.

Using the same simulated admixed populations, we then employed GA-I, CGF-I and HI as core models for performing inference (Figs 3 and S2-S11). With HI, GA, or CGF considered as the true model, our estimation of the optimal model remained highly accurate. On the other hand, when the true model was GA-I or CGF-I, the failure rate decreased from 100% to 30%, compared to the estimation in the previous setting. Furthermore, the estimated time intervals were wider than those of the true ones, although the findings were still more accurate than those using GA and CGF as core models (Table 2).

**Fig. 3:**
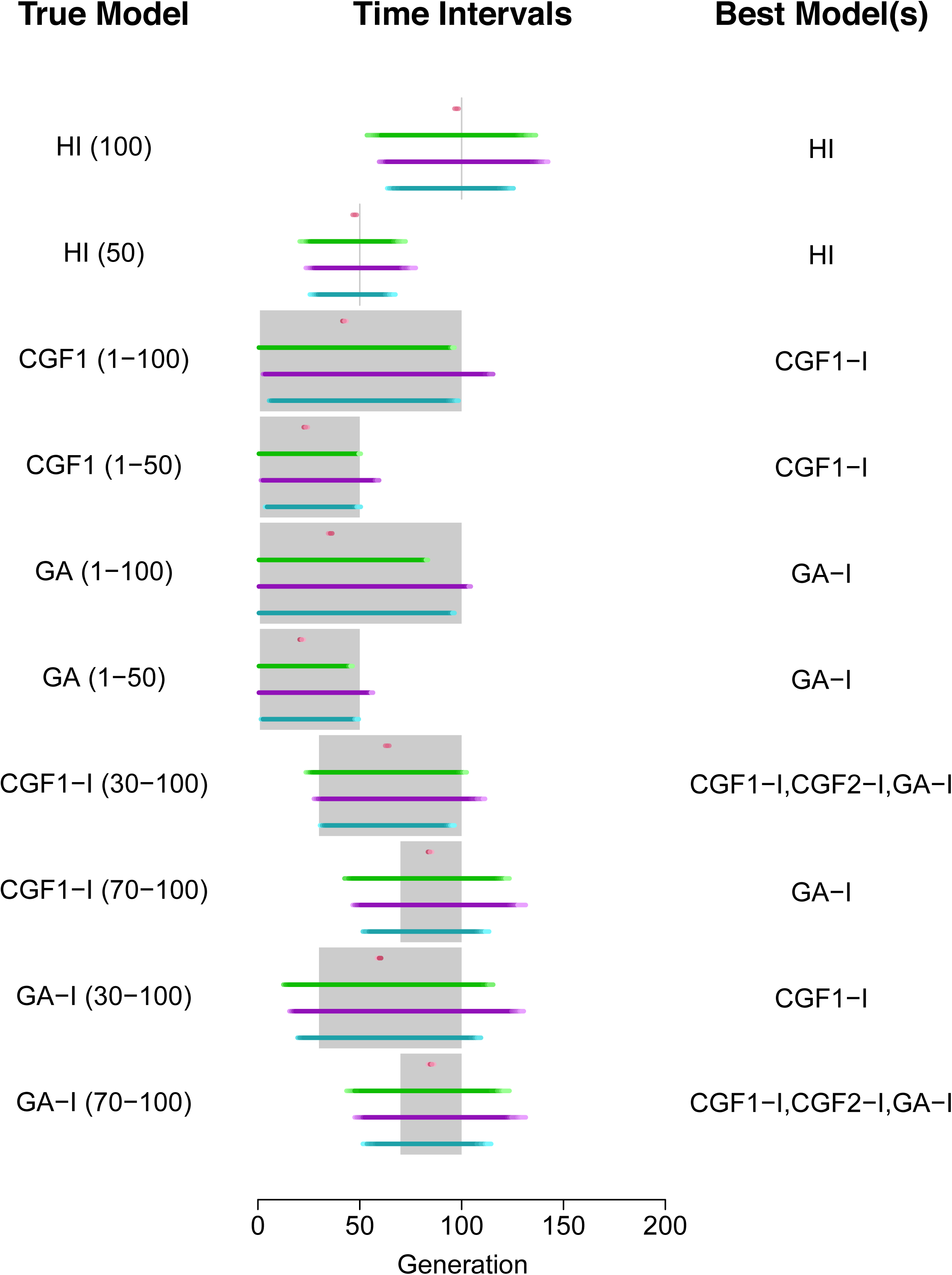
Evaluation of CAMer under various simulated admixture models. Here, the core models are HI, GA-I, CGF1-I, and CGF2-I. The simulated models (True Model) are listed on the left, with the admixture time interval depicted in the parentheses. The gray area on the middle vertical panel is the simulated time interval, whereas colored lines indicate the estimated time intervals under different core models. HI: pink; CGF1-I: green; CGF2-I: purple; GA-I: blue.

### Empirical analysis

We applied CAMer to the selected admixed populations from HapMap, HGDP, and 1KG. For each target population, we first used MALDmef to calculate the weighted LD and fitted the weighted LD with hundreds of exponential functions (Zhou *et al*. 2015). Next, with the weighted LD of target populations, we determined the admixture model and estimated admixture time with CAMer. Quasi F and msE are designed for evaluating the inference with CAMer. The value of msE usually indicates data quality: small msE may indicate a high signal-to-noise ratio (SNR) and vice versa. The quasi F value measures the goodness of fit of the model we employed to fit the admixture event. A small *F* value indicates that the model we used was of satisfactory performance in modeling an admixture event. In our analysis, we used 10^−5^ as the threshold for msE and 1.5 as the threshold for *F*. Therefore, when the msE value ≤10^−5^ and the F value ≤ 1.5, we could not “reject the null hypothesis” that the related model was the true model, i.e., the model well fitted the admixture event. On the other hand, an msE value ≥ 10^−5^ indicates low-quality data that is incapable of identifying the best-fit model, whereas an *F* value ≥ 1.5 prompts us to “reject the null hypothesis” and conclude that the model did not well fit the admixture. In the case of the same population from different databases, the data with smaller msE values were given more credits. For example, we obtained samples of ASW from the HapMap and the 1KG. With the ASW data from HapMap, the best-fit model was HI of 6 generations, and both msE and F values indicated that the inference was acceptable (Fig S12). However, using the ASW data from 1KG, the best-fit models were CGF2-I of 1 to 9 generations and GA-I of 1 to 9 generations (Fig S13). A quasi F value of 2.12 indicated that neither the CGF2-I nor GA-I model satisfactorily fitted the admixture event. Because the msE value of the data set from 1KG was smaller, the conclusion using ASW was as follows: based on the best data we had, the time intervals estimated under the HI, GA-I, CGF1-I, and CGF2-I model were 6 generations, 1–9 generations, 1–13 generations, and 1–9 generations, respectively. Furthermore, none of these models satisfactorily modeled the admixture, whereas the HI model and continuous models (CGF2-I and GA-I) showed better performance. We also applied CAMer to other admixed populations (Table 3, Figs S14–17). MEX was satisfactorily modeled by the CGF model, with the estimated admixture time interval being 1–17 or 1–18 generations, although the identification of the recipient population could not be determined. We also analyzed Eurasian populations, which showed that the Uygurs most likely fit a CGF model with Europeans as donors, with a gene flow lasting for 66 generations to the present. However, the value of msE was larger than 10^2212;^, indicating that the results were not so reliable. The Hazara population experienced a GA-I-like admixture event that lasted for about 5*θ* generations, which started 63 generations ago and ended approximately 5 generations ago.

**Table 3:** Results of CAMer on empirical populations

| Population | Best-fit model | End time | Start time | msE | Quasi.F |
| --- | --- | --- | --- | --- | --- |
| ASW-HapMap (57) | HI | NA | 6 | $2.72 \times 10^{-6}$ | 1.19 |
| ASW-1KG (56) | CGF2-I | 1 | 9 | $1.83 \times 10^{-6}$ | <u>2.12</u> |
| | GA-I | 1 | 9 | $1.86 \times 10^{-6}$ | <u>2.12</u> |
| MEX (86) | CGF1-I | 1 | 17 | $3.56 \times 10^{-6}$ | 1.13 |
| | CGF2-I | 1 | 18 | $3.57 \times 10^{-6}$ | 1.14 |
| MKK (143) | GA-I | 1 | 17 | <u>$1.97 \times 10^{-5}$</u> | 9.82 |
| UIG (10) | CGF1-I | 1 | 66 | <u>$4.01 \times 10^{-5}$</u> | 1.08 |
| Hazara (24) | GA-I | 5 | 63 | $8.53 \times 10^{-6}$ | 1.30 |

2 Number of individuals listed in the parentheses. Values colored in red do not 3 pass our threshold. The time interval is summarized from 22 jackknives, which is 4 shared by more than half of all estimated intervals for continuous models or the 5 nearest integer to the mean of estimated time point for HI model.

## Discussion

Modeling the demographic history of an admixed population and estimating time points of this particular event are essential components of evolutionary and medical research studies (Zhu *et al*. 2004; Zhu and Cooper 2007; Gravel 2012; Jin et al. 2012, 2013; Ni et al. 2015; Zhou *et al*. 2015). Previous methods have employed the length distribution of ancestral tracts (Gravel 2012; Jin *et al*. 2012, 2013), which highly depends on the result of local ancestral inference and haplotype phasing. Another limitation of earlier methods is that only HI, GA, and CGF models were utilized to fit the admixture as well as in identifying the best-fit model. In the present study, our simulations showed that when the true model was not HI, GA, or CGF, the generated inferences were relatively difficult to interpret.

Our method, CAMer, can be utilized in inferring admixture histories by using weighted LD, which can be calculated using genotype data with MALDmef (Zhou *et al*. 2015). Furthermore, we extended the GA and CGF models to the GA-I and CGF-I models in order to infer the time interval for a period of continuous admixture events followed by isolation. Even though CAMer was not consistently very accurate in determining the admixture model, its time interval estimations were reliable.

Two quantities, namely msE and quasi F, were used to evaluate the models inference. These two quantities should both be taken into consideration to identify the best-fit model(s) or the models well fit the admixture. Both the data quality and the goodness of fitting of models can affect the value of msE, although the *F* value mainly measures the goodness of modeling. Therefore, for convenience of interpretation, msE is considered to reveal the data quality and *F* value is considered to evaluate the performance of the model. In our analysis, we suggested thresholds for msE and *F* to determine whether the null hypothesis should be reject or not, which may be too strict in empirical analysis. Actually, both msE and *F* values measure whether the observed weighted LD can be well fitted by the best-fit model(s). For example, the fitting process showed poor performance in the MKK population, which was accompanied by exaggerated msE and *F* values, indicating significant inconsistencies between the observed and fitted weight LD curves (Fig S17). Therefore, in empirical analysis, the msE value reflects the quality of the data, whereas *F* value describes the performance of the model.

In our previous study (Zhou *et al*. 2015), we fitted the weighted LD with hundreds of exponential functions. However, this approach did not fully reveal the occurrence of continuous admixture. To address this issue, the present study developed CAMer to model admixture as a continuous process. We have observed that CAMer performed better than previous admixture model classification methods because it does not require conducting inferences using ancestry tracts, and thus can deal with missing genotypes in the data. CAMer also employed extensions of the classic continuous models, GA-I and CGF-I, which proved to be more flexible in modeling population admixture.

Taken together, CAMer is a powerful method to model a continuous population admixture, which in turn would help us elucidate the complex demographic history of population admixture.

## Software

Our algorithm has been implemented in an R package (R Core Team 2014), named CAMer (Continuous Admixture Modeler). The package is available on the website of population genetic group: http://www.picb.ac.cn/PGG/resource.php or on Github: https://www.github.com/david94048/CAMer

## Author contributions

Conceived and designed the study: **SX**. Developed methods and computer tools: **YZ HQ**. Analyzed the data: **YZ** and **HQ**. Interpreted the data and wrote the paper: **SX YZ HQ**.

## Acknowledgements

None.

